# EEG-ChiMamba: Towards a Robust Mamba-Based Architecture for Dementia Detection from Resting State Electroencephalography

**DOI:** 10.64898/2026.03.23.713697

**Authors:** Christopher Neves, Christopher Steele, Yiming Xiao

## Abstract

Resting-state electroencephalography (rs-EEG) offers a cost effective and portable alternative to conventional neuroimaging for dementia screening, yet the lengthy, multichannel nature of rs-EEG makes learning robust representations challenging. Convolutional and Transformer based architectures dominate current deep learning based approaches, but often struggle with long-range dependencies and may not properly preserve channel-dependent features. In this work, we propose **EEG-ChiMamba**, a state space model based architecture designed for the classification of mild cognitive impairment (MCI) and dementia from normal controls using raw channel-independent rs-EEG signals. Our method decouples channel-wise representation learning from modeling cross-channel interactions and leverages Mamba layers for effective long-sequence modeling. We evaluate our method on the Chung-Ang University EEG dataset (CAUEEG) with 1,155 subjects, the largest public rs-EEG dataset for challenging MCI and dementia differential diagnosis. We achieve a 3-class accuracy of 57.65% using a strict subject-wise split, and relate task-specific features learned by our model as revealed by feature occlusion-based explainability techniques to clinical literature, highlighting that state space models can facilitate interpretable and scalable clinical rs-EEG screening tools for cognitive degeneration. The code for the study is publicly available at: https://github.com/HealthX-Lab/EEG-ChiMamba

## 1 Introduction

Dementia affects more than 58 million individuals globally (Benyumiza et al., 2023) and encapsulates several neurodegenerative diseases, of which Alzheimer’s disease (AD) is the most common. The chronic condition can manifest through a progressive deterioration of cognitive abilities, along with drastic psychological changes, and more than 60% of those affected live in low to middle-income countries (Benyumiza et al., 2023). Individuals commonly show signs of mild cognitive impairment (MCI) before being diagnosed with dementia, with symptom progression that varies depending on the underlying cause. Many treatments and interventions that may slow the course of the disease need an early diagnosis, ideally while a patient is still in the mild cognitive impairment phase or earlier (Jelic & Kowalski, 2009). Biomarkers pointing to dementia-related structural and functional changes in the brain can be used to chart its progression, and structural changes in both macro- and micro-levels for brain morphology and tissue properties, respectively can be identified using Magnetic Resonance Imaging (MRI). Functional MRI (fMRI) and Positron Emission Tomography (PET) are used to study functional and metabolic changes brought on by the disease as potential biomarkers, and recent work has shown that cerebrospinal fluid (CSF) and blood samples can also help monitor the physiological processes underlying Alzheimer’s disease (Schöll et al., 2024). However, most of these potential tests are prohibitively expensive for many, considering that 60% of affected individuals reside in lower-income nations. They are also potentially invasive (e.g., CSF test and PET scans) and all lack portability, making them challenging to administer to remote and/or underprivileged communities. With a rapidly aging population, there exists an urgent need for practical, inexpensive, and accessible diagnostic methods that can offer objective diagnosis and prognosis of dementia.

Electroencephalography (EEG) is a functional imaging alternative that positions itself favorably thanks to its low cost, portability, non-invasiveness, and high temporal resolution. EEG has already shown promise in characterizing functional anomalies arising from the structural changes caused by dementia (Moretti et al., 2007). In addition, irregular EEG patterns have been observed to be common to early-onset dementia arising from many causes, and these irregularities become markedly more distinct in early-onset Alzheimer’s patients (Malek et al., 2017; Micanovic & Pal, 2014), making it a promising tool for an early diagnosis. Compared with typical task-based EEG recording, which needs more elaborate experimental designs and setup, resting-state EEG (rs-EEG) is recorded while a participant is at rest and does not require any complex experimental protocols. This is more convenient than task-based EEG acquisition for both the patients and the EEG technicians, and a technique that can automatically detect dementia from rs-EEG can be of great value for an early and accessible diagnosis.

To date, a wide range of EEG-based biomarkers have been explored to help characterize dementia among the population. Modir et al., 2023 show that the onset of dementia leads to a slowing of EEG dynamics. Measuring the latency of specific Event-Related Potentials (ERP) shows differences between the MCI and AD populations (Santos Toural et al., 2021; J. Zhang et al., 2021). In the spectral domain, studies have linked increases in rs-EEG theta band-power to AD (Farina et al., 2020). Furthermore, a loss of EEG complexity, which characterizes the regularity and predictability of signals (Lau et al., 2022), has also been used to differentiate MCI (Hsiao et al., 2021) and AD (Gaubert et al., 2021) individuals from healthy controls (HCs). However, manual feature extraction often requires extensive preprocessing, and a standardized pipelines are not often used (Craik et al., 2019). More recently, deep learning algorithms have garnered attention as they can automatically learn discriminative features from raw EEG signals without the need for complex preprocessing, which may also have adverse impacts on the downstream analyses (Delorme, 2023). Thanks to the rapid developments in deep learning approaches, developing automated diagnostic tools for the identification of dementia using raw EEG signals, particularly at its early stages, has become a real possibility.

The majority of the existing deep learning solutions applied to the task of AD and MCI detection involve Convolutional Neural Networks (CNN) either applied to raw EEG signals (Ieracitano et al., 2020), two-dimensional spectrograms (Ieracitano et al., 2019) or selective frequency spectrum features (Kongwudhikunakorn et al., 2024). Recurrent models, such as Long-Short Term Memory (LSTM) and Recurrent Neural Networks (RNN) that were designed to process sequential data, have also been used to classify both raw EEG signals (Alvi et al., 2022) and hand-crafted features (Alessandrini et al., 2023) to some degree of success, and Transformer models, which model contextual importance of tokens in sequential data with self-attention, have also achieved promising results when being applied to 2D spectral representations (Miltiadous et al., 2023) and raw signals (Jiang et al., 2024) for EEG. However, learning salient features from raw rs-EEG data is much more challenging, as the lack of apparent signal responses to external event-based stimuli means that deep learning techniques must instead capture sophisticated symptom-related hidden characteristics in the recordings. In this regard, recurrent DL architectures like LSTMs become difficult to train on the lengths of data seen in rs-EEG studies due to their limited memory capacity and non-parallelizable nature, while Transformers often struggle to model complex features unless given large amounts of training samples, which can be unrealistic for EEG experiments. They also suffer from a computational complexity that scales quadratically with input length due to their self-attention mechanism, making it difficult for them to train on long signal sequences without additional innovation. Therefore, convolutional neural networks are often strong choices for these tasks, as they are more robust to these issues and can be competitive with other methods (Kiessner et al., 2024; M.-j. Kim et al., 2023). Additionally, the validity of some existing methods on how temporal convolutional networks and Transformers have been applied to time-series data has recently been challenged (Luo & Wang, 2024; Zeng et al., 2023). Traditionally, deep learning techniques jointly embed the input channels of a multivariate time series. More specifically, as traditional deep learning models process the multivariate time series, they mix all input channels simultaneously during a projection to a higher dimensional feature space. This means that each feature learned by the model will contain information from all input channels (Nie et al., 2023). However, recent work has shown that better outcomes can be achieved by treating each separate input channel as a univariate time series and projecting each one to a separate feature space. This implies that each feature will contain learned patterns from only one input channel. It is believed that the increase in performance resulting from the channel-independent modeling approach may also be due to the observation that greater differences exist between channels in a multivariate time series like EEG than in computer vision tasks, where only the Red-Green-Blue channels are present (Luo & Wang, 2024). Others have argued that methods combining all input channels *simultaneously* fail in multivariate time-series tasks because they assume that each input channel contains data emanating from the same underlying process (Han et al., 2024). This assumption does not hold true in EEG, where although electrodes may share some signal components due to effects like volume conduction, they ultimately capture the activity of many distinct underlying neural processes (Cohen, 2017). This univariate modeling strategy has found some success in EEG but is still not widely adopted, and some method of learning relationships between separate input channels can still be beneficial. Notably, some recent EEG DL methods have benefited from the univariate modeling strategy, ranging from EEG data synthesis with generative diffusion DL models (Vetter et al., 2024) to more accurate seizure detection and classification (Tang et al., 2023). Integrating these insights and finding more effective ways of dealing with very long sequences is crucial to being able to model lengthy rs-EEG sequences and capturing far-reaching physiological signatures in the data.

Very recently, state space models (SSM) have positioned themselves as a strong option for very long sequence modeling. The SSM framework describes the behavior of a dynamical system by modeling it as a collection of states and how the system transitions between these states. Deep state space models, such as Mamba (Gu & Dao, 2023), have achieved state-of-the-art performance in challenging long-range sequence tasks with results that match and often exceed Transformer models while boasting a computational complexity that scales linearly with sequence length. This enhanced scalability, in contrast to Transformers, is beneficial when handling the high sampling rates of EEG data, but what an effective Mamba-based DL model for robust EEG feature extraction looks like remains an open question. For the task of sleep stage and sleep disorder classification, Siddhad et al., 2024 propose a dual-branch Mamba architecture, and C. Zhang et al., 2024 combine a bi-directional Mamba with attention. In terms of multi-task EEG classification, Gui et al., 2024 propose a bi-directional Mamba architecture with a task-aware mixture of experts to perform epilepsy, sleep stage, emotion, and motor-imagery classification. Behrouz and Hashemi, 2024 design a hybrid Mamba and graph neural network model capable of processing EEG and fMRI data, and Panchavati et al., 2024 modified a U-Net architecture with Mamba layers for seizure detection. For motor-imagery classification, Yang and Jia, 2024 apply Mamba across both the temporal and channel dimensions to extract relevant EEG features. To the best of our knowledge, the approach by Tran et al., 2024 is the only other Mamba-based method for EEG-based differential diagnosis for dementia. Specifically, they attempt to detect Alzheimer’s disease and frontotemporal dementia from the resting state EEG of 88 participants. However, they perform a trial-wise validation in their experiments (subjects may have data present in both training and testing splits), which may severely overestimate model performance as it trivializes individual differences between subjects and degrades generalization (Brookshire et al., 2024).

In this work, we intend to address the aforementioned issues with a deep learning method EEG-ChiMamba, applied to rs-EEG with the following contributions. **First**, we design a novel Mamba-based DL model to address the need for long-range sequential modeling techniques in rs-EEG signal classification, which we use to allow differential diagnosis of dementia (i.e., HC vs. MCI vs. dementia classification). Specifically, our method uses a channel-independent modeling approach with effective temporal and channel mixing strategies to extract robust EEG features. **Second**, we are the first to benchmark a Mamba-based architecture using the first large-scale dementia rs-EEG dataset (M.-j. Kim et al., 2023), and show improved classification performance over existing methods while using substantially fewer parameters. **Third**, by using occlusion-based explainability methods, we examine the features learned by the proposed DL model and reveal key physiologically relevant insights regarding dementia and mild cognitive decline.

## 2 Materials and Methods

Figure 1 outlines our proposed DL architecture for HC, MCI and dementia classification based on rs-EEG. *First*, input EEG samples are reshaped so that each electrode channel (referred to as **channel** throughout the text) can be treated as a univariate time series throughout the model. *Second*, we apply patching to parse the input time series into discrete segments of length *P* while independently projecting each channel to a *D*-dimensional **feature** vector. Next, signals are processed by 3 EEG-ChiMamba blocks. Each block after the first is preceded by a dropout layer and a fully connected layer that projects the number of features per electrode to the model dimension (Dropout + Project). Within each EEG-ChiMamba block, a channel-wise LayerNorm and a Mamba SSM layer learn relationships between sequence segments. Then, a channel mixer captures interactions between the *C* number of channels. *Finally*, outputs from the last EEG-ChiMamba block are average-pooled and concatenated with the age of the participant to obtain a final vector of shape *X* ∈ R^(^*^C^*^×^*^D^*^)+1^, which is then classified using a linear layer as HC, MCI, or dementia. To mitigate overfitting to the training dataset, we incorporate Dropout layers (Srivastava et al., 2014) for regularization. We apply Dropout before the fully connected layer in the Dropout + Project block and in all three of the EEG-ChiMamba blocks on the outputs of the Mamba SSM.

**Figure 1:**
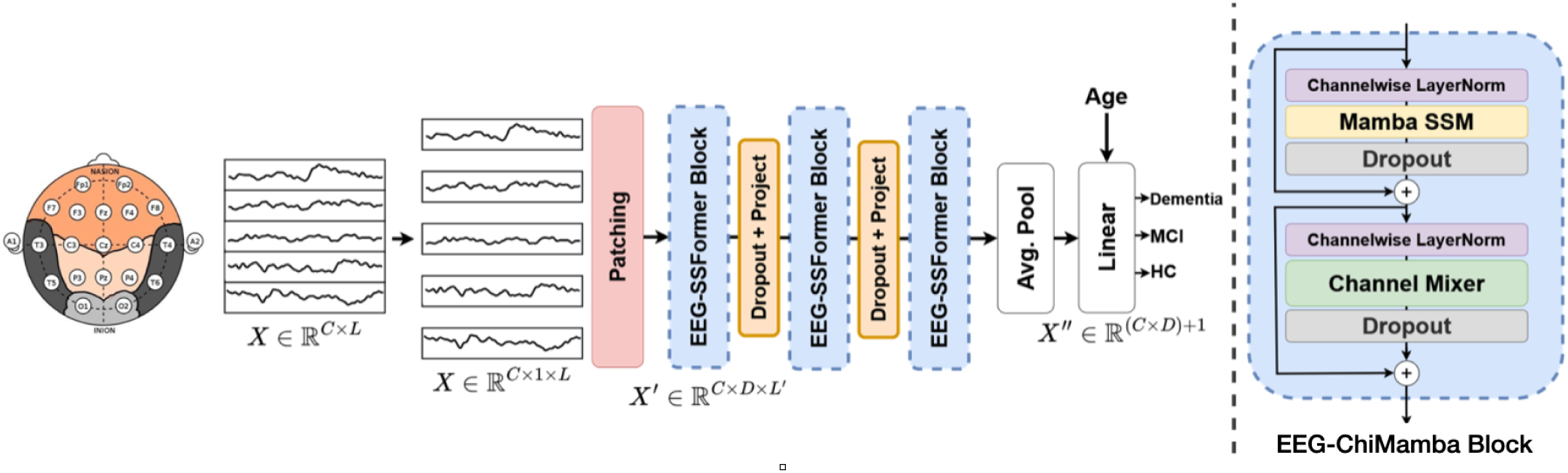
Overview of the model architecture for HC, MCI, and Dementia classification. *C, D, L* represent the sizes of the channel, feature, and temporal dimensions.

In the following sections, we will provide an overview of the different components of our proposed DL model, including the time-series patching, channel-independent feature extraction, and channel mixing. We then describe the dataset used, experimental setup, and techniques employed to interpret model outputs for potential physiological insights.

### 2.1 Channel independent feature learning

We adopt the channel-independent feature modeling strategy employed by previous works (Liu et al., 2024; Luo & Wang, 2024; Nie et al., 2023) to better capture distinct electrode-specific features in the input EEG data and reduce the effects of distribution shift (Han et al., 2024). This is done by applying the patching, projection, channel-wise LayerNorm, and Mamba operations separately for each channel. Cross-channel interactions are captured by a separate channel mixing step (Channel Mixer in Fig. 1).

#### 2.1.1 Time series patching

Before passing through the model, the input EEG signal undergoes z-score normalization using the training set’s mean and standard deviation. When using the age signal, the mean and standard deviation of the training set subject’s age is used.

We then perform patching on the input signals. The input time series is parsed into smaller segments of *P* number of time steps, where *P* is referred to as the patch length. These segments of time series data are then projected to a higher dimensional feature space of dimension *D*. Normally, this patching operation (parsing + projection) is performed in two distinct steps (Nie et al., 2023). However, we perform patching using a single 1D convolution layer, similar to the work of Luo and Wang, 2024. Specifically, the fully convolutional patching first reshapes input signals to *X* ∈ R*^C^*^×1×^*^L^*. Next, we apply a 1D convolution with a stride and kernel length equal to the patch size *P* . The convolution operation will result in a patched output *X*^′^ ∈ R*^C^*^×^*^D^*^×^*^L′^*, where *L*^′^ is equal to the number of segments. Unlike individual words in Natural Language Processing tasks, a single time step does not have semantic meaning or context. Patching extracts local semantic information between groups of time steps and reduces the overall computational complexity of the model (Nie et al., 2023), motivating its use in our DL architecture. After the patching operation, the input data is normalized in a channel-wise manner using an inverted LayerNorm technique, which will be described in the following section.

#### 2.1.2 Inverted LayerNorm

Unlike standard z-score normalization that is applied to input data at the preprocessing stage, LayerNorm is a mini-batch processing technique that is applied to normalize processed intermediate outputs between layers. The commonly used LayerNorm module can have two main issues. First, when normalizing all features for a single time step, the resulting time series will contain few variations between features, reducing the representation power. Second, due to the large differences between time series channels, large fluctuations related to an event in one channel may introduce spurious noise in another when using the standard LayerNorm technique, thus removing the benefits of a channel-independent modeling strategy. To solve these, the inverted LayerNorm technique was introduced by Liu et al., 2024 and normalizes data across the time steps rather than the features. This channel-wise normalization has been shown to be more robust to distribution shifts and more effective when dealing with non-stationary signals (T. Kim et al., 2021; Liu et al., 2022, 2024). The detailed formulation of the inverted LayerNorm is described in Equation 1, where *µ_n_* and *σ_n_* are the mean and standard deviation of the timeseries of Feature *D* from Channel *C*.

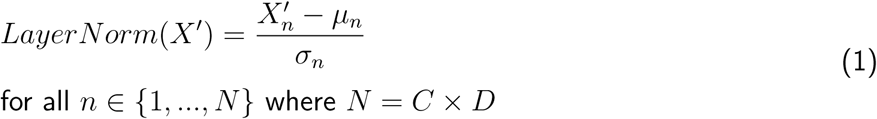

#### 2.1.3 Learning temporal dependencies using Selective State Spaces

After the *inverted LayerNorm*, we use a Mamba state space model (SSM) to extract global temporal relationships between time steps from the data. State space models describe dynamic systems and project an input signal *x*(*t*) to a hidden state *h*(*t*), which is then used to obtain an output state *y*(*t*). This is performed through Equation 2, where the *M_A_* matrix governs how the hidden state *h*(*t*) changes over time, *M_B_* decides how the current input affects the hidden state, *M_C_* maps the state to the output, and *M_D_* allows the input to directly modulate the output.

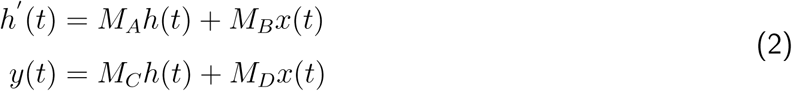

Mamba improves upon previous state space models like S4 (Gu et al., 2022) by allowing the parameters *M_B_* and *M_C_* to vary based on the input using a selective scanning algorithm. This allows Mamba to filter irrelevant portions of an input sequence while highlighting important information in a data-dependent way. This filtering process requires input channels to be mixed and projected to a higher-dimensional space, so we reshape inputs so that Mamba is applied independently for each input channel *C*. In this work, we take advantage of Mamba’s powerful long-range sequential modeling to learn global temporal patterns in rs-EEG data.

### 2.2 Decoupled channel-and-feature mixing

The patching layer, linear projection layers and Mamba SSM jointly learn both local and global temporal patterns while treating each channel as a separate univariate time series. An important next step is to capture important cross-feature and cross-channel relationships. In this work, similar to other timeseries works (Liu et al., 2024; Luo & Wang, 2024), we decouple the channel and feature mixing steps into two separate operations.

Many approaches learn feature and channel relationships in a coupled approach. That is, they learn inter-channel and inter-feature interactions jointly in a single mixing step, often using a simple convolution (M.-j. Kim et al., 2023). However, when using channel-independent modeling approaches, this coupled channel and feature modeling can substantially increase parameter counts. Decoupling the two operations not only drastically reduces the parameter count of the model which decreases computational complexity, but may also force the model to use the parameters more efficiently (Luo & Wang, 2024).

In our work, feature mixing is performed on a per-channel basis in the patching layer, the linear projection layers (Dropout + Project in Fig. 1) and the Mamba SSM. Then, to capture cross-channel relationships, the data is reshaped and permuted to *X*^′^ ∈ R^(^*^D^*^×^*^C^*^)×^*^L′^* and we perform two point-wise grouped convolutions with a number of groups equal to the number of feature dimensions per electrode channel, *D*.

In our final computational framework, we employed the channel mixing in the spatial domain. However, alternative spectral domain mixers have been shown to be effective, and we further validate our decision to perform spatial domain-based mixing against the spectral domain counterpart for the designated application. We present this comparison as an ablation experiment in Section 2.2.2.

#### 2.2.1 Channel mixing in the spatial domain

We implement the channel mixing step using a series of point-wise convolutions in the spatial domain. A point-wise convolution is a convolution that uses a kernel size of 1 and is functionally similar to a linear layer. We use two point-wise convolutions following an inverted bottleneck structure. The first point-wise convolution projects the number of channels *C* to be twice as wide as the input dimension, making the inputs go from *X*^′^ ∈ R^(^*^D^*^×^*^C^*^)×^*^L′^* to *X*^′^ ∈ R^(^*^D^*^×2^*^C^*^)×^*^L′^* . Afterwards, we apply a ReLU non-linearity, followed by a second point-wise convolution, to return the number of channels back to their original dimension.

#### 2.2.2 Channel mixing in the spectral domain

Recent work has shown that frequency domain mixers may sometimes lead to better modeling results (Patro & Agneeswaran, 2024; Yi et al., 2024) for time-series data compared to their spatial counterparts. To that end, we test if this is also the case for rs-EEG classification.

To model the relationships between channels in the spectral domain, we adapt the frequency domain mixer initially proposed by Patro and Agneeswaran, 2024 called EinFFT. First, data that has been patched and processed by the Mamba SSM is reshaped and permuted to *X*^′^ ∈ R*^B^*^×^*^L′^*^×(^*^D^*^×^*^C^*^)^. The EinFFT mixer then performs the Fourier transform of the processed input data to obtain X ^′^ ∈ C*^B^*^×^*^L′^*^×(^*^D^*^×^*^C^*^)^. The data is then linearly transformed using complex-valued weights W and biases B, which essentially acts as a linear layer in the frequency domain. A ReLU nonlinearity is then applied to the transformed output, followed by a second linear transformation in the frequency domain, after which the signals are finally converted back to the spatial domain using the inverse Fourier transform.

In their work, Patro and Agneeswaran, 2024 parameterize the complex weight matrix W as a block diagonal matrix to reduce the overall parameter count of the model. They set a fixed number of 4 blocks in their parametrization, meaning that data will be mixed using the linear layers only among elements in the same block. To enable the channel mixing mentioned above, we remove the fixed constraint on the number of blocks in W and instead set the number of blocks to *D* (corresponding to the number of features per channel). In their work, Patro and Agneeswaran, 2024 convert the inputs to the frequency domain by performing the 2D Fast Fourier Transform (FFT) over the sequence and block dimension. We maintain this implementation, and apply the FFT across the equivalent (*L*^′^*, D*) dimension of the processed input data. Setting the number of blocks to *D* emulates the grouped convolution mentioned in Section 2.2, where the non-linear transformation is applied only to the channel dimension. It is important to note that both the spatial and spectral mixers have a similar number of learnable parameters to allow for a fair comparison.

### 2.3 Dataset and preprocessing

For our study, we used the Chung-Ang University Hospital EEG (CAUEEG) dataset (M.-j. Kim et al., 2023), which is the largest rs-EEG dataset of patients with various stages of dementia to date. Our experiments focus on the dementia subset of this dataset, which categorizes subjects as healthy controls (HC), having mild cognitive impairment (MCI), or diagnosed with dementia. This dataset includes rs-EEG data of 1,155 subjects, with some recorded during photic stimulation. The dataset is subdivided into training, validation, and testing sets. The training dataset contains 950 subjects (367 HC, 334 MCI, and 249 dementia), the validation set contains 119 subjects (46 HC, 42 MCI, and 31 dementia), and the testing set contains 118 subjects (46 HC, 41 MCI, 31 dementia). Most participants have well-annotated clinical diagnoses of dementia subtype, and the sex distribution consists of 6 males for every 10 females.

Note that the sex of each specific participant in the dataset is removed to preserve their anonymity. The means and standard deviations of the ages for all HC, MCI, and dementia subjects in the training, validation, and testing datasets are detailed in Table 1.

**Table 1:** Mean and standard deviation of the ages (years) for all subjects in the training, validation and testing dataset splits.

| Condition | Training | Validation | Testing |
| --- | --- | --- | --- |
| Healthy Controls | 65.37±9.48 | 64.91±10.43 | 63.21±8.43 |
| Mild Cognitive Impairment | 73.71±7.83 | 75.02±7.13 | 72.75±8.57 |
| Dementia | 76.59±8.01 | 77.25±9.57 | 77.25±7.17 |

In order to be classified as HC, MCI, or dementia, M.-j. Kim et al., 2023 use a series of inclusion criteria, which we summarize here. The criteria for a healthy control include: **1)** no interruption in daily activities **2)** results on neuropsychological tests were normal (within a standard deviation of age and education-adjusted baselines) (Ahn et al., 2010; Jahng et al., 2015). To be classified as having MCI, the following criteria must be met: **1)** no interruption in daily activities **2)** there must have been complaints regarding issues with memory **3)** neuropsychological tests show cognitive impairment (impairment must be ≥ 1 standard deviation of age and education adjusted norms) (Ahn et al., 2010; Jahng et al., 2015) **4)** 0.5 rating in clinical dementia **5)** the subject is not categorized as demented according to the DSM-IV criteria (First & Pincus, 2002). Finally, patients with dementia are identified if they follow the criteria outlined by the National Institute of Neurological and Communicative Disorders and Stroke and Alzheimer’s Disease and Related Disorders Association, as well as the DSM-IV (Dubois et al., 2007). We will refer the readers to the original publication (M.-j. Kim et al., 2023) for further details.

For the curated data, each subject has a minimum of 5 minutes of EEG recordings, sampled at 200 Hz and recorded using 19 EEG electrodes placed according to the 10-20 placement system. An EKG or ECG electrode and a channel for the photic stimulus are also included. Since this study aims to explore the classification capabilities of deep learning architectures on raw rs-EEG data, the photic stimulation and EKG/ECG channels are left unused in our case. The EEG data is band-pass filtered between 0.5 and 70 Hz at the time of acquisition and is referenced to the common average. Following M.-j. Kim et al., 2023, we do not perform any further pre-processing, and provide models with 10 seconds of EEG data that are randomly sampled from each participant’s total available data after ignoring the first 10 seconds of each recording (to mitigate setup artifacts). This study is the first to apply a Mamba-based architecture to this corpus, and the size of the dataset ensures a robust evaluation of model performance.

### 2.4 Experimental setup and ablation studies

To assess the effectiveness of our proposed methods, we conduct experiments comparing them to various baselines and ablated configurations in terms of HC vs. MCI vs. dementia classification performance. To measure classification performance, we compute the macro-averaged classification accuracy, macro-averaged AUROC, and the class-wise F1-scores for all baselines and model configurations. We calculate each metric over 5 random seeds and report each metric’s mean and standard deviation.

As CNNs are the most commonly used DL architectures in EEG classification and the best-performing models tested by M.-j. Kim et al., 2023 on the CAUEEG dataset were CNNs, we implement the strongest CNN models from the previous investigation of M.-j. Kim et al., 2023 as baselines. These include a 1D-ResNet-18 and the best performing 1D-VGG model (1D-VGG-19). We use the model hyperparameters determined by M.-j. Kim et al., 2023 for the CNN models.

In addition, we perform experiments to validate various design choices for our proposed DL architecture using the validation set subjects of the CAUEEG corpus. **First**, we assess the optimal domain for channel mixing by comparing our model using the spatial channel mixer described in Section 2.2.1 (EEG-ChiMamba-PW) against one model variant using its spectral counterpart (EEG-ChiMamba-EinFFT). **Second**, we quantify the effectiveness of the inverted LayerNorm technique described in Section 2.1.2 on EEG signals by testing a variant of our model with the conventional LayerNorm (EEG-ChiMamba-PW w/o I-LN). **Finally**, since age has been shown to be a critical risk factor for dementia (McCullagh et al., 2001), we integrate it into our prediction pipeline similar to other works (M.-j. Kim et al., 2023). We name this model EEG-ChiMamba-PW + Age. This model is compared with a model variant without utilizing such information (EEG-ChiMamba-PW).

All models are trained using the same random-cropping scheme as in (M.-j. Kim et al., 2023), where a training sample consists of 10 seconds of signal randomly cropped from a subject’s data. This acts as a form of regularization and helps models see a larger variety of signals. We discard the first 10 seconds of each participant’s data to avoid recording artifacts emanating from the trial start. During training, each model sees 100,000 random crops of signals per epoch over a total of 50 training epochs, resulting in 5,000,000 random crops seen per model over their training regime. We use identical training, validation, and testing splits as (M.-j. Kim et al., 2023), who split data in a subject-wise manner to avoid data leakage for fair evaluation. For the validation and testing sets, we split each participant’s data into deterministic (non-randomly sampled) 10-second non-overlapping windows, resulting in 9123 samples in the validation set (3153 HC, 3435 MCI, 2535 dementia) and 8795 samples in the testing set (3027 HC, 3350 MCI, 2418 dementia). All models are trained using the AdamW optimizer (Loshchilov & Hutter, 2017). For the SSFormer models, we use a base learning rate of 0.0001 and a minibatch size of 32, and a cosine decay learning rate scheduler. The 1D-ResNet-18 and 1D-VGG-19 models are trained using the hyperparameters outlined in (M.-j. Kim et al., 2023). That is, the 1D-ResNet-18 model uses a base learning rate of 0.000469 and a minibatch size of 512, along with a cosine learning rate scheduler. The 1D-VGG-19 model is trained using a base learning rate of 0.000469 and a minibatch size of 256. We apply the same level Gaussian noise as a data augmentation to the input signals for all methods to improve robustness of model training.

The EEG-ChiMamba architecture in Fig. 1 employs 3 EEG-ChiMamba blocks with a hidden feature size of 128 per channel. We use a dropout rate of 0.2 applied to the Mamba SSM outputs in all EEG-ChiMamba blocks, and a rate of 0.05 for the channel mixers. We use a patch size of 8 and a patch dimension of 64 for all model variants. For the experiment involving age, we first normalize the age value using the mean and standard deviation of the training set, and add small amounts of random Gaussian noise to the value. This prevents the model from memorizing the age of the subject, and overfitting to the subjects in the training dataset and is also performed by M.-j. Kim et al., 2023. We then concatenate it to the average pooled results before classifying them using Linear layers.

### 2.5 Model interpretability

Beyond the designated classification task, investigation of the discriminative features crucial to the task can also offer relevant clinical knowledge to better understand the diseases. Therefore, we probe our model for physiologically relevant insights using an occlusion sensitivity analysis on both the individual EEG channel electrodes and each of the canonical frequency bands.

#### 2.5.1 Channel occlusion sensitivity topographic maps

We generate channel occlusion sensitivity topographic maps to understand which EEG electrodes/channels (or scalp regions) are the most important for the downstream HC vs. MCI vs. dementia classification task.

We first select the subset of correctly classified HC, MCI, and dementia samples from the testing set subjects using the EEG-ChiMamba-PW model variant. We then sequentially occlude each electrode by setting its value to 0, then measure the associated drop in predicted class probability from our DL model for the correct class, and record the average value across all samples in the relevant cohort. Note that the class probabilities are obtained by applying the softmax function to the model’s output logits. These channel-wise probability changes are then mapped over an outline of a scalp according to their positions in the 10-20 EEG electrode placement system, and intermediate values between the electrode positions are interpolated using cubic interpolation with the MNE-Python library (Gramfort et al., 2013; Larson et al., 2025) in order to generate a smooth heatmap over the scalp surface. The values in the resulting topographic maps further from 0 imply that the signals from that electrode were more important to the model’s final classification. A positive number represents a drop in predicted class probability, whereas a negative number signifies that the probability for the predicted class increased after occluding the electrode. The latter may happen if those channels are particularly noisy or do not contain useful information for the predicted class and only serve to confound the model. Since the topographic maps are generated using correctly classified samples, electrodes that drop the class-specific probability better reflect the importance of the electrode for determining that class. The occlusion sensitivity topographic maps are generated and averaged across 5 random seeds.

#### 2.5.2 Canonical frequency band analysis

To understand which canonical frequency bands are more relevant to the classification task with our model, we iteratively band-stop filter each of the delta (0.5-4 Hz), theta (4-8 Hz), alpha (8-13 Hz), beta (13-30 Hz), and gamma (30-90 Hz) bands of test dataset as specified by (Van Diessen et al., 2015). For each occluded frequency band, we calculate the new classification accuracy of the model. We then compare this new classification accuracy with the accuracy obtained without removing frequency bands and compute the **relative** change. We accumulate this relative accuracy change over 5 random seeds using the EEG-ChiMamba-PW + Age model and report the mean and standard error. If the model made use of features mostly present within a certain frequency band, then we expect a large drop in relative accuracy once that specific band is removed.

## 3 Results

### 3.1 Classification performance of baseline models and ablation studies

Table 2 presents the classification accuracies of the proposed model configurations on the validation split of the CAUEEG dataset. The best performing model based on the accuracy and AUROC on the validation set is then selected and evaluated alongside the included CNN baseline models on the held-out testing split of the dataset. On the validation set, the channel mixing in the spatial domain (EEG-ChiMamba-PW, 59.27 ± 0.93%) outperforms the model variant with mixing in the spectral domain (EEG-ChiMamba-EinFFT, 53.99 ± 1.31%). When switching the LayerNorm module to perform normalization across tokens (EEG-ChiMamba-PW) instead of across features (EEG-ChiMamba-PW w/o I-LN), we see a rise in all classification metrics with a 2.48% rise in classification accuracy.The model configuration that includes age (EEG-ChiMamba-PW + Age) performs the strongest among all EEG-ChiMamba architectures on the validation set. This configuration includes the token-wise LayerNorm and spatial channel mixing.

**Table 2:** Validation set results for EEG-ChiMamba configurations. Best results are in bold, second best results are underlined.

| Method | Acc. (%) | Macro AUROC | HC F1 | MCI F1 | Dementia F1 |
| --- | --- | --- | --- | --- | --- |
| EEG-ChiMamba-EinFFT | 53.99±1.31 | 0.727±0.010 | 0.575±0.006 | 0.475±0.017 | 0.582±0.022 |
| EEG-ChiMamba-PW w/o I-LN | 56.79±1.94 | 0.759±0.017 | 0.594±0.009 | 0.505±0.022 | 0.626±0.039 |
| EEG-ChiMamba-PW | 59.27±0.93 | 0.775±0.005 | 0.617±0.011 | 0.533±0.004 | 0.651±0.017 |
| **EEG-ChiMamba-PW + Age** | **61.47±1.32** | **0.798±0.012** | **0.641±0.017** | **0.558±0.012** | **0.669±0.019** |

The test set results are outlined in table 3. With an accuracy of 57.65 ± 0.67%, our proposed method (EEG-ChiMamba-PW) outperforms all of the CNN baselines and model variants on the test set. With the inclusion of the age signal, the performance jumps to 58.37 ± 0.391% (EEG-ChiMamba-PW + Age). Compared to the CNN baseline models, we confirm the effectiveness of the Mamba-based architecture. EEG-ChiMamba-PW outperforms both the 1D-ResNet-18 and 1D-VGG-19 models on average. All models perform the best on HC classification on the test set, and the performance of the MCI group is lowest across all methods despite the MCI group having a greater number of training subjects in comparison to the dementia group (334 vs. 249, respectively).

**Table 3:** Classification results of baseline models and Final EEG-ChiMamba configurations on the Test Set. Best results are in bold, second best results are underlined.

| Method | Acc. (%) | Macro AUROC | HC F1 | MCI F1 | Dementia F1 |
| --- | --- | --- | --- | --- | --- |
| 1D-ResNet-18 | 51.88±1.27 | 0.693±0.011 | 0.642±0.012 | 0.414±0.011 | 0.491±0.025 |
| 1D-VGG-19 | 54.01±1.04 | 0.712±0.013 | 0.668±0.008 | 0.448±0.012 | 0.497±0.020 |
| EEG-ChiMamba-PW | 57.65±0.67 | 0.760±0.006 | 0.706±0.005 | 0.484±0.009 | 0.513±0.013 |
| **EEG-ChiMamba-PW + Age** | **58.37±0.39** | **0.769±0.001** | **0.723±0.005** | **0.482±0.011** | **0.521±0.018** |

In terms of parameter counts, the 1D-VGG-19 model is the largest with 20.2 million parameters, followed by the 1D-ResNet-18 model with 11.3 million parameters. The EEG-ChiMamba-PW, EEG-ChiMamba-EinFFT and EEG-ChiMamba-PW + Age models have approximately 3.8 million parameters.

### 3.2 Channel occlusion sensitivity topographic maps

Figure 2 shows the channel occlusion scores described in Section 2.5.1. The channel occlusion scores are mapped to a birds-eye view of the scalp surface using their corresponding positions in the 10-20 placement system.

**Figure 2:**
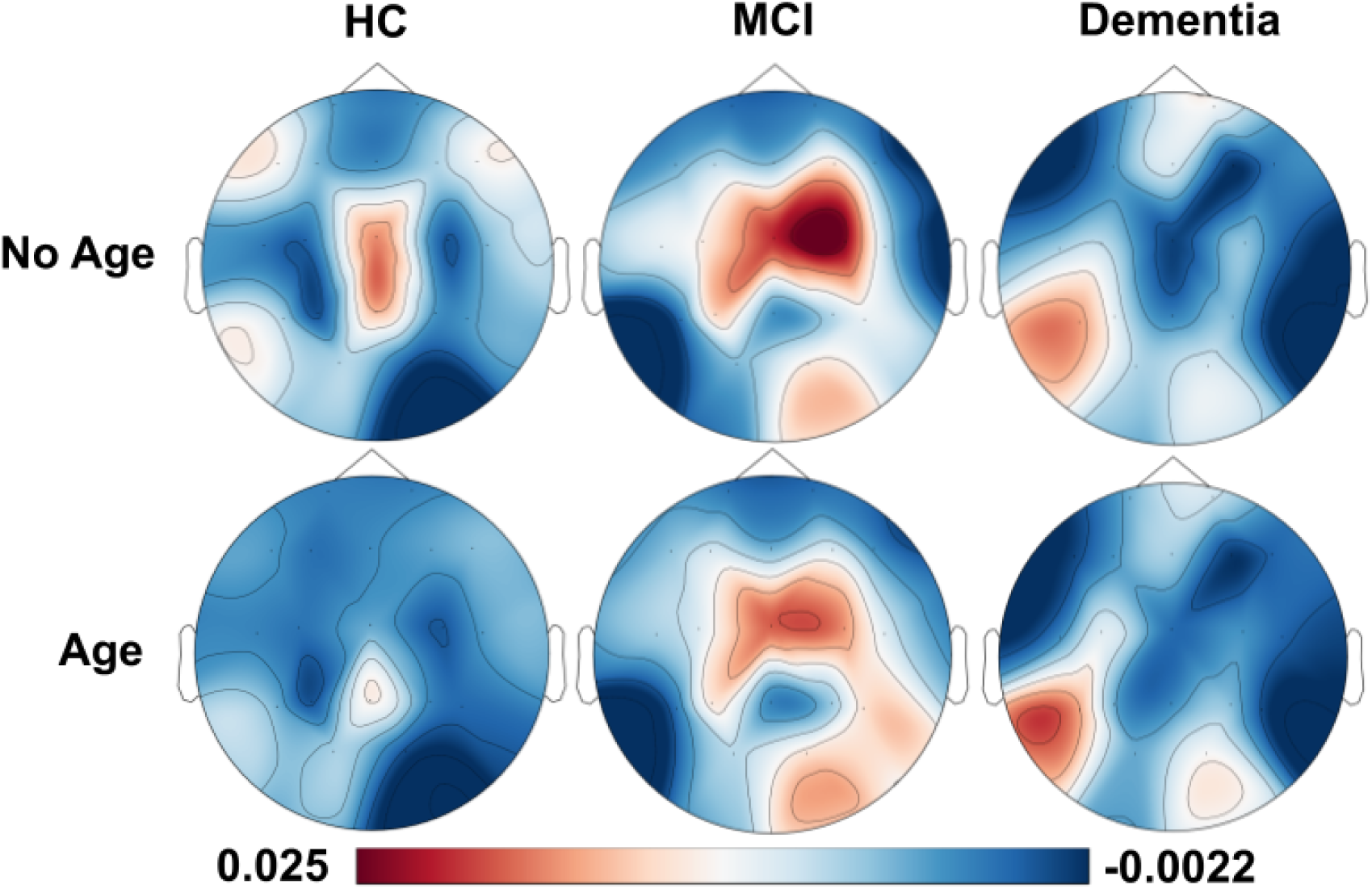
Group-wise channel occlusion maps for EEG-ChiMamba model trained without and with the age signal.

The EEG-ChiMamba-PW and EEG-ChiMamba-PW + Age models are used to obtain predictions from subjects in the test set. The correctly predicted samples aggregated over 5 random seeds are used to generate the topographic maps.

The top row of the figure shows the occlusion scores for the EEG-ChiMamba-PW model trained without the age signal. The bottom row depicts the results for the EEG-ChiMamba-PW + Age model, which includes the age signal. There are smaller fluctuations in class probability when using a participant’s age, whereas the model without age is more prone to larger prediction probability changes when zeroing out a channel. This can be observed by the smaller range of magnitudes for the occlusions with age.

The electrodes that have the greatest effect on predicted probability are over the frontal lobe, left-temporal and central-parietal lobe for HC, the central-parietal, right-temporal, and occipital lobe for MCI, and mostly over the occipital and temporal-left lobe for dementia subjects. The inclusion of age leaves these areas relatively unchanged across conditions but increases the importance of the inion of the scalp for the MCI group.

### 3.3 Canonical frequency band analysis

The relative changes in class accuracies after iteratively band-stop filtering each canonical frequency band and the corresponding standard error of the mean calculated over 5 random seeds are shown in Figure 3.

**Figure 3:**
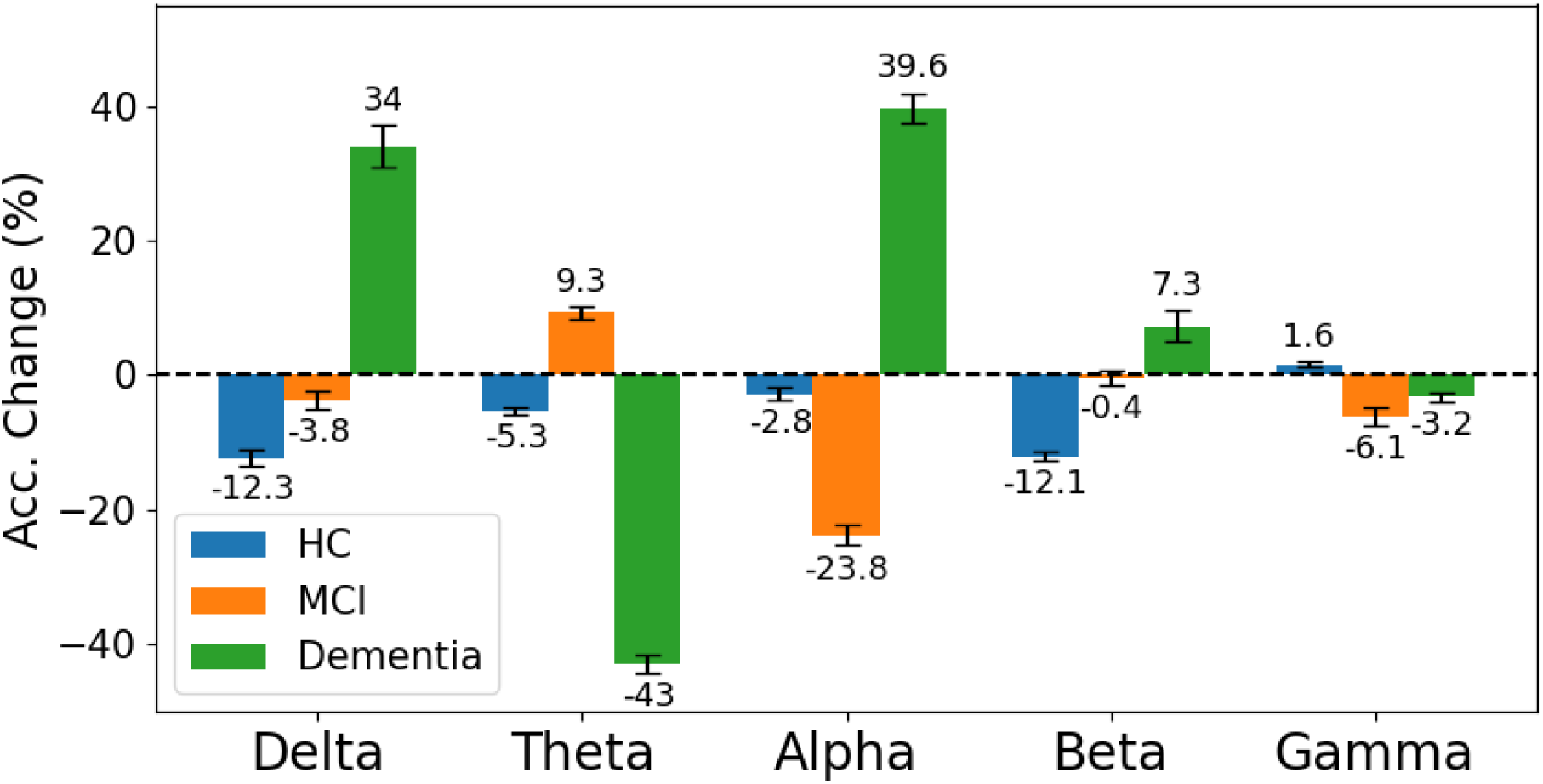
Relative accuracy change of best performing EEG-ChiMamba model configuration with canonical frequency band-stop filter.

Removing the delta band has the highest relative accuracy decrease for the HC class with a 12.3% drop in classification accuracy compared to the non-occluded performance. It also results in a substantial performance gain for the Dementia group, with a 34% increase in relative accuracy. The removal of the theta band yields the greatest drop in classification performance for the Dementia group, reducing the classification accuracy by 43% whereas ignoring the alpha band increases the performance on the Dementia group by 39.6%. This provides the highest performance increase for the Dementia group. However, removing the Alpha band also drops the performance on the MCI samples by 23.8%, making it the steepest performance decrease for the MCI group across frequency occlusions. HC performance drops to a level comparable to the removal of the Delta band when occluding Beta frequencies, and has a small effect on the MCI and Dementia groups. The Gamma band occlusion yields the smallest change in accuracy for the HC and Dementia groups, with a modest drop of 6.1% for MCI samples.

## 4 Discussion

In this work, we present a novel Mamba-based channel-independent architecture that extracts salient features from raw rs-EEG signals for classifying dementia while outperforming models with more than four times as many parameters. This suggests that Mamba-based architectures may be more suitable than pure CNN architectures, which have achieved excellent results in EEG classification tasks. Additionally, our model is validated on the largest public dementia rs-EEG databases, and exploring the features learned by our model reveals physiologically relevant insights.

While gaining strong popularity in computer vision and natural language processing domains, Mamba has seldom been used for rs-EEG classification and even less for dementia detection tasks. Recently, Tran et al., 2024 use an ensemble model featuring Mamba to classify inputs containing both raw EEG and manually extracted spectral features for the task of discriminating between subjects with Alzheimer’s disease, frontotemporal dementia, and healthy individuals. To the best of our knowledge, they are the only existing work attempting to apply Mamba to dementia detection using EEG, but their use of manually extracted features steer the scope of the task away from using minimally processed signals. In addition, they use a trial-based experimental protocol, where they shuffle the extracted EEG segments from all participants and split the segments into training and testing sets. As a result, in this setup, a model will be able to see data from a single subject in both the training and testing splits (i.e., data leakage), allowing the model to memorize subject-specific instead of task-specific details, reducing the algorithm’s generalizability to unseen subjects. Other studies have applied more traditional DL techniques for dementia classification. (Sen et al., 2024) use the intrinsic time-scale decomposition to extract rotation components from EEG signals, then use a 1D CNN to classify patients with Alzheimer’s disease in an in-house dataset. However, they also employ the trial-based validation setup and report a classification accuracy of 94%. (Radwan et al., 2024) use graph neural networks with Granger causality graphs for the binary classification of abnormal EEGs with the CAUEEG dataset (M.-j. Kim et al., 2023). (Paillard et al., 2025) integrate neuroscientific priors into the design of a lightweight DL architecture that employs learnable wavelets and Riemannian geometry to classify EEG signals. They achieve strong results on a pre-processed version of the CAUEEG dataset, and make use of the phase-locking value feature to capture the effects of the photic stimulation that some subjects were exposed to during the signal recording. Finally, (Farina et al., 2020) compare machine learning classifiers trained separately on manually extracted features from fMRI and rs-EEG data for AD vs. HC, MCI vs. HC, and AD vs. MCI classification. Similar to the results reported in Table 3, they also show that the MCI group is the hardest to classify correctly out of the three conditions. Due to the difference in algorithm validation setup and differences in the datasets, it is difficult to directly compare the classification accuracy of our method against these aforementioned works.

To validate our approach, we compare our work to the strongest individual models developed by M.-j. Kim et al., 2023, which include the popular ResNet-18 and VGG-19 CNN architectures adapted for EEG data and tested on the CAUEEG dataset. Our spatial EEG-ChiMamba variant (EEG-ChiMamba-PW) with inverted layernorm achieves a 3.64% improvement over the next best-performing baseline model (1D-VGG-19) without the inclusion of the age signal. Notably, our proposed architecture contains 3.8 million trainable parameters, substantially less than the 20.2 million parameters of the 1D-VGG-19 and the 11.3 million parameters of the 1D-ResNet-18, suggesting that using a state space model and a channel-independent approach allows for learning the long-range features unique to each channel that are discriminative for the downstream classification task more efficiently.

In the ablation experiment comparing the mixing of the model’s hidden features and input channels in the spatial and spectral domains, the spatial mixing achieves superior classification scores on the validation set, outperforming the spectral mixing by 5.28%. Although the spatial mixer was selected as the final architecture based on superior validation set performance, we tested the spectral mixer (EEG-ChiMamba-EinFFT) on the test set in a post-hoc exploratory analysis. While this variant achieved comparable test set performance, it showed substantially lower validation set performance along with greater variability across random seeds suggesting less stable generalization behavior. This highlights an important challenge in model selection using EEG data: architectural variants may not perfectly preserve their rankings across dataset splits due to the inherent variability of neurophysiological data. This is further exacerbated in a subject-wise dataset splitting regime.

The frequency mixer may not have been as effective as the spatial mixer in our application as it applies the Fourier transform across all elements of the sequence and feature dimensions (*L, D*, which means that it observes the global periodic components of the data but fails to localize important frequency components in time. On the other hand, time-frequency techniques such as the short-time Fourier transform are so valuable in EEG data as they combine the best of both worlds and are capable of determining *when* an important change in signal frequency occurs (Morales & Bowers, 2022). By minimizing the temporal component of the data, the spectral channel mixer may be ignoring important characteristics in the data that could lead to a correct dementia classification.

Although frequency domain variate mixing (including the EinFFT layer) has shown success in time-series tasks before (Patro & Agneeswaran, 2024; Patro & Agneeswaran, 2025; Yi et al., 2024), the datasets used contain a maximum input length of 336 time steps. This number is dwarfed by the 2000 time steps used for 10 seconds of rs-EEG data in our experiments, and modeling the variations in frequency components over time may be less important. In addition, Patro and Agneeswaran, 2024; Patro and Agneeswaran, 2025; Yi et al., 2024 test the frequency domain mixing for time-series forecasting tasks, which may be better suited to this form of channel mixing.

To better understand the decision-making process of our deep learning algorithm, we investigate the contributions that individual EEG channels and frequency bands have on the classification outcomes. In classic EEG analysis, these factors are often of great interest as potential biomarkers for disease diagnosis or insights to better understand the target neural processes. In the channel occlusion plots in Figure 2, we observed similar patterns of channel importance between the models with and without using the age factor in the classification. Overall, the model that makes use of age shows lower changes in prediction probability, especially in the healthy control population, suggesting that the introduction of the age signal can lead to greater robustness in the classifier. This is likely due to the fact that advanced aging is often correlated with cognitive function decline (Sanford, 2017), and the age signal adds confidence to the models identification of HC.

Notably, for MCI, our model attributes greater importance to central, frontocentral, right parietal and right occipital areas of the scalp, which is consistent with findings by (Chetty et al., 2024) showing that the left-frontal, frontocentral, central and parietal areas exhibit significantly higher gamma/alpha ratios in the prodromal AD (MCI) participants when compared to healthy controls. In terms of the dementia subjects, there is a lower emphasis on central regions of the scalp as shown in Figure 2, and a higher emphasis is placed on occipital regions, with some important electrodes located on the left posterior regions. Tabbal et al., 2025 study the deviations in relative spectral power of each scalp electrode from normative models in HC, AD and subjects with Parkinsons disease. They show that individuals with AD show deviations in some occipital channels in the theta band, and left temporal parietal channels at the beta band. This overlaps with the electrodes in 2 that show decreases in prediction probability when occluded.

We also investigate the importance of features learned throughout individual canonical frequency bands on the diagnostic task. As shown in Figure 3 the theta frequency band is crucial for dementia detection. This is consistent with the conclusion of Farina et al., 2020 that theta power is the strongest predictor of Alzheimer’s disease status in resting-state EEG, and they report that the significance of the theta band is consistent across multiple regions of the scalp. Chetty et al., 2024 come to the same conclusion when discriminating between Alzheimer’s disease, prodromal Alzheimer’s disease, and healthy controls from resting-state EEG features. They find that the theta power of Alzheimer’s patients was elevated compared to their prodromal and healthy counterparts. Besides theta band, previous studies Giustiniani et al., 2023; Kudo et al., 2024 also suggested increased delta band in rs-EEG to be associated with Alzheimer’s disease. In our case, the removal of the delta band contributed to a large increase in the classification accuracy of dementia. This may be due to the fact that we focus on the symptom of dementia across a number of underlying causes other than specifically AD. Tabbal et al., 2025 also identify a correlation between delta band power and scores obtained on the Mini-Mental State Examination (MMSE) in patients with Alzheimer’s disease. It is possible that the large relative increases in performance of the dementia group that were observed in the removal of the delta and alpha bands and the corresponding drops in relative accuracy for the HC and MCI subjects may be due to the elimination of confounding signals that led to the misclassifications of dementia patients, but were still valuable for discriminating between HC and MCI.

It is important to note that there is substantial heterogeneity between individuals even among clinical groups, and those with MCI can have large amounts of overlap in neurophysiological characteristics with other conditions (Tabbal et al., 2025). For healthy controls, we find the most discriminative frequency band lies in the delta and beta bands. This finding is mirrored in the work of Farina et al., 2020, who state that beta band power is the most significant predictor of healthy control status. Finally, Chetty et al., 2024 report increased gamma vs. alpha power ratio and gamma-band functional connectivity in the prodromal Alheimer’s disease population, which may help explain the drop in relative classification accuracy that our model experiences for the MCI class when removing the alpha and gamma bands.

Although we show improved performance using our novel DL technique, some limitations to the current work remain that can provide opportunities for future explorations. The CAUEEG dataset is the largest rs-EEG dataset for dementia, but there exists a few sources of heterogeneity among subjects that may impact our classification accuracy. First, the dataset contains two different experimental protocols, with a sub-cohort of patients receiving photic stimulation during data acquisition. Although our model was still capable of learning relevant EEG features for differential diagnosis, the discrepancy in experimental protocols may interfere with the learning process in some cases, or can be taken advantage of by implementing manually extracted features like the phase-locking value to increase classification performance (Paillard et al., 2025). Another source of heterogeneity lies in the diversity of disease diagnoses among the population in the dataset. While the majority of the subjects in the dataset were assigned general labels of dementia, MCI, or HC, the causes of dementia vary (e.g., Parkinson’s disease dementia vs. vascular dementia), which may contribute to different characteristics in EEG activities. Future investigations that reveal the common and distinct EEG patterns of different causes for MCI and dementia may help better reveal the mechanisms of the symptoms.

## 5 Conclusion

In this work, we propose a novel Mamba-based channel-independent DL model for HC vs. MCI vs. dementia classification. Using a subject-wise validation scheme, we develop and test our proposed method based on the CAUEEG dataset with the largest corpus of rs-EEG collected from dementia patients. Our results demonstrate superior performance compared to strong CNN models while reducing the model parameter count by approximately four times. Furthermore, we show that our model is capable of extracting physiologically relevant features from the resting state signals, with insights in line with the current neuroimaging literature. Our results offer a promising avenue to leverage rs-EEG for the diagnosis and study of dementia, which is particularly beneficial for remote areas and underprivileged communities.

## Data and Code Availability

The code is publicly available at: https://github.com/HealthX-Lab/EEG-ChiMamba.

